# The unique protein-to-protein carotenoid transfer mechanism

**DOI:** 10.1101/127183

**Authors:** E.G. Maksimov, N.N. Sluchanko, Y.B. Slonimskiy, K.S. Mironov, K.E. Klementiev, M. Moldenhauer, T. Friedrich, D.A. Los, V.Z. Paschenko, A.B. Rubin

## Abstract

List of abbreviations

OCP: Orange Carotenoid Protein
NTD: N-terminal domain of OCP
CTD: C-terminal domain of OCP
COCP: C-terminal OCP-related Carotenoid-binding Protein
COCP-W288A: COCP with a Ala substitution of Trp
GFP: enhanced Green Fluorescence Protein
FRP: Fluorescence Recovery Protein
NTE: N-terminal extension (comprising the αA helix up to amino acid 20)
ΔNTE: OCP with the 20 most N-terminal amino acids deleted (OCP^Δ2-20^)
ECN: Echinenone
CAN: Canthaxanthin
AL: Actinic Light

**Abstract:** Orange Carotenoid Protein (OCP) is known to be an effector and regulator of cyanobacterial photoprotection. This 35 kDa water-soluble protein provides specific environment for keto-carotenoids, the excitation of which induced by the absorption of blue-green light causes dramatic but fully reversible rearrangements of the OCP structure, including carotenoid translocation and separation of C- and N-terminal domains upon transition from the basic orange to photoactivated red OCP form. While recent studies significantly improved our understanding of the OCP photocycle and interaction with phycobilisomes and the fluorescence recovery protein, the mechanism of OCP assembly remains unclear. Apparently, this process requires targeted delivery and incorporation of a highly hydrophobic carotenoid molecule into the water-soluble apoprotein of OCP. Recently, we introduced a novel carotenoid carrier protein, COCP, which consists of dimerized C-domain(s) of OCP and can combine with the isolated N-domain to form transient OCP-like species. Here, we demonstrate that *in vitro* COCP efficiently transfers otherwise tightly bound carotenoid to the full-length OCP apoprotein, resulting in formation of the photoactive OCP from completely photoinactive species. We accurately analyze peculiarities of this carotenoid transfer process which, to the best of our knowledge, seems unique, previously uncharacterized protein-to-protein carotenoid transfer process. We hypothesize that a similar OCP assembly can occur *in vivo*, substantiating specific roles of the COCP carotenoid carrier in cyanobacterial photoprotection.

## 1. Introduction

Adapted to perceive the macroworld, human senses constantly fail to acknowledge the most incredible yet ordinary capabilities of biological materials on the nanoscale. Translated into units we can oversee, the absorption of a single green 500-nm photon by a 35-kDa protein compares to the traceless swallowing of a 4 g, 9 mm Parabellum bullet, fired from a shotgun at the speed of sound, by a 35-g slab of soft biological matter. This apocalyptic event, taking place in myriads at every twinkle of the eye, forms the basis of oxygenic photosynthesis for the benefit of every form of Life on the Planet. Whereas the photosystems and their elaborate light-harvesting antennae undertake every effort not to waste incident photons, the sheer chemistry of light-induced reactions at the same time required the development of protective mechanisms against photodamage by reactive oxygen species and other radicals. Cyanobacteria developed a tightly controlled system of photoprotection to shield their photosynthetic reaction centers from the harmful effects of intense solar radiation, in which the 35 kDa Orange Carotenoid Protein (OCP) plays the central role. As if to carry things to extremes, it is the 567 Da carotenoid cofactor 3’-hydroxyechinenone (hECN) performing the task of photon absorption to bring about OCP photoconversion from the basal orange state OCP^O^ to the red active state OCP^R^, the latter being able to tightly interact with the soluble phycobilisome (PBs) antennae complexes to quench their fluorescence and prevent the flow of excessively absorbed excitation energy to the photosystems [1-8].

The structural organization of OCP is modular, with an N-terminal and C-terminal domain (NTD and CTD, respectively) of about equal size, which symmetrically encapsulate one carotenoid molecule within a central cavity [9-11]. Whereas OCP can accommodate a variety of xanthophylls, photo- and PBs quenching activity is only possible with 4-(4’-)ketolated carotenoid derivatives. The importance for the keto moiety is underlined by the fact that the only specific protein-carotenoid interactions exist in the basal OCP^O^ form, comprising two hydrogen bonds from Trp-288 and Tyr-201 in the CTD to the 4-keto oxygen at the terminal β-ring [12-15]. In OCP^O^, the compact structure is further stabilized by protein-protein interactions across the large inter-domain interface including the Arg-155/Glu-244 salt bridge and a contact formed between the short αA-helix in the 20 amino acid-long N-terminal extension of the NTD with a specific part of the outer surface of the CTD [16, 17]. Upon photon absorption and isomerization of the carotenoid, the specific protein-chromophore H-bonds break, the αA-helix detaches and unfolds from the CTD and both domains separate with the carotenoid sliding by about 12 Å completely into the NTD [18-20].

Already the first attempts to purify OCP from native sources revealed a contamination by a red 16 kDa fragment, which was termed Red Carotenoid Protein (RCP) and was later identified as the OCP-NTD coordinating the carotenoid, which might be formed from full-length OCP *in vivo* (and forms *in vitro)* by proteolysis [11, 21, 22]. Deciphering of more and more cyanobacterial genomes revealed that OCP is broadly distributed, but, astonishingly, a large fraction of cyanobacterial genomes harbor genes for one or multiple isolated genes for NTD as well as CTD homologs on top of the OCP gene [23, 24]. Some cyanobacteria even harbor only genes for NTD and CTD homologs. Whereas the functional role(s) of the NTD homologs, which have been termed helical carotenoid proteins or HCPs for their exclusively α-helical secondary structure elements, has well been established as being the effector modules of the photoprotective mechanism, the role and importance of the multiple and wide-spread CTD homologs, though being structurally related to the widespread superfamily of mixed α/β-structured nuclear transport factor-2-like (NTF-2, PFAM 02136) protein domains, was only vaguely defined as “accessory” or “modulatory” since CTD fragments purified from cyanobacteria never contained carotenoids. Only recently, using carotenoid-producing *E. coli* strains for protein production, it was shown that the OCP-CTD (termed COCP for C-terminal OCP-related carotenoid-binding protein) indeed forms a carotenoid-binding entity by employing a symmetric, highly stable dimeric arrangement, in which two COCP monomers coordinate a single, symmetrically oxygenated xanthophyll such as canthaxanthin (CAN) or zeaxanthin (ZEA) [25]. This highly symmetric configuration entailed the by far most red-shifted absorption spectrum from all OCP derivatives characterized so far, very low chirality in the visual CD spectra and distinct, unprecedented features at the characteristic v_1_ to v_4_ bands in Raman spectra [25]. Moreover, upon mixing of COCP(CAN) with the apoprotein form of the NTD (Apo-RCP), spectral and functional features of an OCP^O^-like state appeared, which readily underwent photoconversion resulting in the formation of RCP that was capable of PBs fluorescence quenching [25]. In effect, these findings revealed an as yet unprecedented carotenoid transfer mechanism initiated by COCP suggesting that isolated COCP homologs in cyanobacteria may play some role in carotenoid mobilization and storage, besides acting as singlet oxygen quenchers [23, 25, 26]. The modular architecture of OCP and the possibility to assemble it in fully functional form from its separated domains allows tailoring the use of carotenoid-binding entities according to highly specialized functions needed by the cell.

The processes leading to chromophore integration during maturation of OCP are not known yet. Notably, our previous work indicated that carotenoid transfer from COCP also occurred into the apoprotein of full-length OCP [25]. Here, we study this process in detail to elucidate the reaction steps leading to assembly of functional OCP by spectroscopy and bioanalytical techniques. Carotenoid transfer starts from the violet COCP with an interaction of the N-terminal extension (NTE) of the OCP apoprotein with COCP followed by dissociation of dimeric COCP as a rate-limiting step, thus uncapping the carotenoid for insertion into the NTD of the OCP apoprotein and resulting in a red intermediate that spontaneously underwent conversion to the OCP^O^ state. During the transfer, several reaction intermediates are identified that rely on the intimate interactions existing between isolated CTDs or NTD-CTD, respectively, some of which including mixed OCP dimers connected via a single carotenoid molecule, and these assemblies are characterized in detail by analytical size-exclusion chromatography and small-angle X-ray scattering.

## 2. Materials and methods

### 2.1. Protein cloning, expression and purification

Cloning, expression and purification of the His6-tagged *Synechocystis* OCP^WT^ apoprotein, its apoprotein form lacking the N-terminal extension (NTE) comprising the first 20 amino acids (ΔNTE), and FRP were described previously [13, 14, 16, 27]. The His-tag was identical in all the protein forms used. Holoforms of COCP and COCP-W288A were expressed in echinenone (ECN) and canthaxanthin (CAN)-producing *Escherichia coli* cells essentially as described before [25]. All proteins were purified by immobilized metal-affinity and size-exclusion chromatography to electrophoretic homogeneity and stored at +4 °C in the presence of sodium azide. The apoproteins could be stored frozen at −80 °C.

Cloning and expression of fusion GFP::OCP protein. A cDNA fragment coding for GFP was amplified by PCR with Pfu polymerase using GFPBHI_F (CAC<u>GGATCC</u>GAGCAAGGGCGAGGAGC) and GFPBHI_R primers (GGCCG<u>GGATCC</u>TTGTACAGCTCGTCC) both carrying *BamHI* restriction sites (underlined) using the pGFP plasmid (Clontech Laboratories) as a template. The PCR product was digested with *Bam*HI, agarose-gel purified and cloned in-frame into the *Bam*HI-digested dephosphorylated OCP-pQE81L vector. Verification of plasmids with the required orientation of GFP-OCP cDNA in a expression ORF was performed using PCR with Scrng_F (GGGCATCGACTTCAAGGAGG) and Scrng_R (CGGTGACCAGCTTGCATAGG) primers. Plasmids from PCR-positive clones were tested by sequencing. The *E. coli*-strain carrying appropriate plasmids for carotenoid (ECN/CAN) synthesis for production OCP holoprotein was described earlier [13]. Expression was performed in auto-inducible media [28]. Recombinant GFP-OCP fusion protein was purified on HisTrap HP column (GE Healthcare Life Sciences). Eluted protein was dialyzed against standard PBS solution.

### 2.2. Absorption and fluorescence spectroscopy

Absorption spectra were recorded using a Maya2000 Pro (Ocean Optics, USA) spectrometer as described in [14, 20, 25]. Upon absorption measurements, a blue light-emitting diode (LED) (M455L3, Thorlabs, USA), with a maximum emission at 455 nm was used for the photoconversion of the samples (further termed actinic light (AL) for OCP^O^→OCP^R^ photoconversion). Fluorescence spectra and decay kinetics were recorded by USB4000+ (Ocean Optics, USA) CCD spectrometer and home-build spectrometer based on single photon counting modules (Becker&Hickl, Germany), respectively, as described in [20]. Temperature of the sample was stabilized by a Peltier-controlled cuvette holder Qpod 2e (Quantum Northwest, USA) with a magnetic stirrer.

### 2.3. Analytical size-exclusion chromatography

To study concentration dependences of hydrodynamics of proteins and the interaction between FRP and ΔNTE, we pre-incubated protein samples (100 μL) and subjected them to size-exclusion chromatography (SEC) on a calibrated Superdex 200 Increase 10/300 column (GE Healthcare) equilibrated with a 20 mM Tris-HCl buffer, pH 7.6, containing 150 mM NaCl, 0.1 mM EDTA, and 3 mM β-mercaptoethanol and operated at 25 °C at 1.2 mL/min flow rate. Unless otherwise indicated, the elution profiles were followed by carotenoid-specific absorbance (wavelengths are specified). In some cases, the column was constantly illuminated by a blue-LED to achieve OCP photoconversion. All experiments were performed at least three times using independent preparations of proteins.

### 2.4. Native gel-electrophoresis

The individual COCP (30 μM), Apo-OCP (30 μM), or the mixtures of COCP (30 μM) with increasing concentration of Apo-OCP (from 0 to 140 μM) were incubated for 30 min at 33 °C and then subjected to gel-electrophoresis under non-denaturing conditions at pH 8.6 [29], essentially as described earlier [14]. To reveal changes only in the carotenoid-bound forms of COCP and Apo-OCP, the gels were scanned without any staining.

### 2.5. Chemical crosslinking

Chemical crosslinking of COCP, Apo-OCP, or their mixture by glutaraldehyde (GA) was performed essentially as described previously (REF to BBA and FCS paper). Control samples did not contain GA. After 30 min pre-incubation the protein content was analyzed by SDS-PAGE.

### 2.6. Small-angle X-ray scattering (SAXS) data collection and processing

OCP^O^ and its apoprotein (Apo-OCP) were analyzed by synchrotron SAXS at the P12 beamline (PETRA III, DESY Hamburg, Germany) using a batch mode (OCP^O^) or an inline HPLC system for sample separation immediately preceding data collection (OCP^O^ at low concentration; Apo-OCP at high concentration). To this end, the centrifuged samples were either directly analyzed by SAXS using a sample changer or were loaded in a volume of 75-100 μl on a Superdex 200 Increase 10/300 column (GE Healthcare) pre-equilibrated with filtered and degassed 20 mM Tris-HCl buffer (pH 7.6) containing 150 mM NaCl, 0.1 mM EDTA, 2 mM dithiothreitol, and 3 % glycerol. SAXS curves collected at different OCP^O^ concentrations in batch mode showed substantial concentration dependence, indicating that no extrapolation is possible, and thus two extreme situations (low and high concentration) were analyzed individually. The SAXS curve for OCP^O^ at high concentration (175 μM), collected in batch mode (exposure time - 0.05 s, dead time – 0.05 s, temperature – 20 °C) and having little noise, was used to model oligomerized OCP^O^. For SEC-SAXS analysis, to ensure maximal dimerization and spatial separation from residual monomers, Apo-OCP was loaded in high concentration (200 μM), whereas OCP^O^ was instead loaded in low concentration (38 μM), to eliminate possible dimerization while allowing for collecting the amount of frames sufficient to obtain a moderately noisy SAXS curve upon averaging (as low as 20 μM OCP^O^ concentration measured in batch mode produced a noisy curve of non-satisfactory quality, not suitable for modeling). Chromatography was conducted at 20 °C with a 0.5 ml/min flow rate, and the flow was equally divided between the SAXS and TDA detection modules to ensure simultaneous data collection from equivalent parts of a profile. TDA allowed simultaneous analysis of the eluate by absorption at 280 nm, refractive index (RI), and right angle light scattering (RALS). The RALS data for OCP samples and those for the bovine serum albumin (BSA) standard were used to obtain Mw distribution for OCP peaks (dn/dc was taken as 0.185). SAXS data frames (exposure time – 1 s, dead time – 1 s, λ=1.24 Å) for the buffer (500 first similar frames) and the sample (frames 1517-1542 for Apo-OCP and frames 1750-1820 for OCP^O^) were collected. No radiation damage was detected by inspection of the time course of the scattering for protein frames. All indicated buffer frames were averaged and subtracted from each protein frame. Protein frames in the indicated ranges were then scaled to the curve corresponding to the peak maximum and averaged by PRIMUS [30] to produce the resulting SAXS curve to be utilized for modeling using the ATSAS package (http://www.embl-hamburg.de/biosaxs/software.html). Guinier regions of all SAXS curves analyzed were linear, did not reveal any signs of interparticle interactions and were used to determine experimental R_g_ values. P(r) distributions calculated by GNOM [31] at S ≤ 0.25 Å^-1^ were used to estimate D_max_ values. *Ab initio* molecular envelopes of OCP and Apo-OCP at either low or high protein concentrations were built by running 20 DAMMIF calculations followed by DAMAVER averaging [32, 33]. Theoretical SAXS curves and fitting of the crystallographic OCP^O^ monomer (PDB ID 4XB5) or dimer (PDB ID 3MG1) to the experimental data were calculated using Crysol [34]. In order to model the structure of the Apo-OCP dimer, the CTD-CTD dimeric core (residues 185-317 from the 4XB5 structure) was fixed according to the proposed symmetric COCP structure [25], whereas the NTDs (residues 3-161 from the 4XB5 structure) were allowed to move freely and independently, while being connected to CTDs by natural flexible linkers (residues 162-184). The N-terminal hexahistidine tag and its connection to the NTD (17 residues overall) were also considered flexible in both chains. Such a structure (P1 overall symmetry) was modeled using CORAL [35] by minimization of the discrepancy between the model and the experimental SAXS curve calculated by Crysol.

## 3. Results and discussion

### 3.1. Absorption spectroscopy reveals the carotenoid transfer between COCP and Apo-OCP

As shown in our recent study, when expressed in carotenoid-producing strains of *E. coli*, the CTDs of OCP can effectively form rather stable protein dimers symmetrically linked by a single carotenoid molecule (COCP), and mixing COCP with Apo-RCP produced a transient OCP^O^-like species [25]. This indicated that a unique carotenoid transfer between the OCP domains takes place, however, the transient OCP^O^-like species were very unstable and eventually developed into a differently colored species that was spectrally and functionally indistinguishable from holo-RCP [25].

Surprisingly, replacement of Apo-RCP by the full-length Apo-OCP in the carotenoid transfer experiment presents significant advancement. Indeed, upon addition of colorless Apo-OCP to a violet solution of COCP, the absorption of the sample changes dramatically and, in equilibrium, resembles the absorption of the orange OCP form with its characteristic vibronic structure with multiple peaks (Figure 1A). The resulting spectrum can be decomposed into the sum of spectra from an orange (OCP^O^) and violet (COCP) forms (dashed and dotted lines in Figure 1A, respectively), indicating that, under the conditions used, 73% of the carotenoid is transferred from COCP into OCP, while 27% remain in some form that is spectrally indistinguishable from COCP. The extent of the OCP^O^ state formation depended on the concentration of Apo-OCP used, and the effect, evaluated as a decrease in absorbance at 550 nm (see Figure 1B and D) or as a concomitant increase in absorbance at 470 nm (data not shown), saturated at approximately one Apo-OCP per one COCP dimer ratio (Figure 1C). However, under all conditions tested, even at high Apo-OCP to COCP concentration ratios, the remaining spectral contribution from the violet COCP-like forms never vanished completely. This observation is striking, especially since the carotenoid transfer occurred spontaneously, which should result in vanishing of the initial violet COCP species assuming that the backward transfer of the carotenoid from OCP to Apo-COCP does not occur, as determined in independent experiments [25]. But, as we will outline below, this feature of the carotenoid transfer results in more than one product.

**Figure 1.**
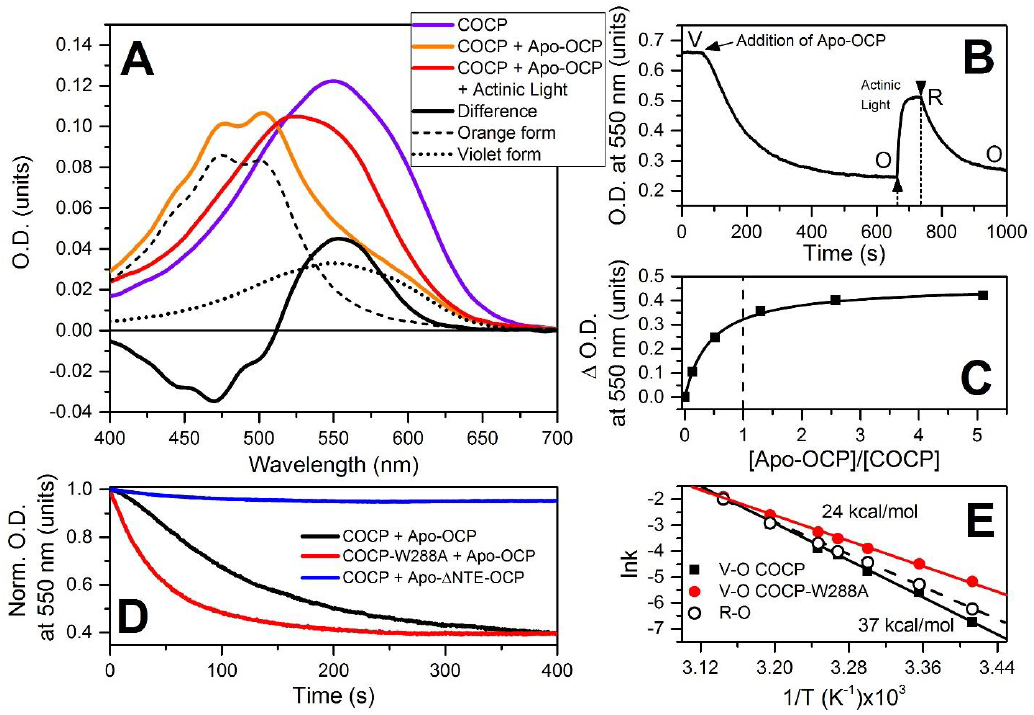
Carotenoid transfer from COCP to Apo-OCP followed by absorption spectroscopy. **(A)** – absorption spectrum and color of the carotenoid bound to COCP undergo significant changes upon addition of Apo-OCP (3 Apo-OCP per 1 COCP dimer), which gradually turns the sample from violet into red and, finally, into orange. The spectrum of species obtained after mixing of COCP and an excess of Apo-OCP represents a mixture of two forms – orange (dashed line), reminiscent of the OCP^O^ spectrum, and violet (dotted line), similar to the absorption spectrum of initial COCP solution. **(B)** – characteristic time-course of O.D. at 550 nm measured upon addition of Apo-OCP to the solution of carotenoid-containing COCP (V – violet) resulting in formation of a typical orange-like OCP form (O), which is photoactive and could be converted in to the red form (R). **(C)** – changes of O.D. at 100 seconds after addition of different Apo-OCP concentrations to the solution of COCP dimer at 33 °C. **(D)** – time-courses of carotenoid shuttling upon addition of Apo-OCP to COCP (black), Apo-OCP to COCP-W288A (red) and Apo-ΔNTE-OCP to COCP (blue). **(E)** – Arrhenius plots of the rates of the orange form formation for COCP and COCP-W288A as the initial sources of carotenoid. Dependences were approximated by linear functions to estimate the activation energies.

Thus, conversion of the violet form into the orange one (V-O) occurs spontaneously, albeit relatively slowly, upon mixing of COCP and Apo-OCP, suggesting that the hydrophobic carotenoid is passed directly between the proteins evading the solvent. Interestingly, for such a process to take place, the COCP dimers obviously must dissociate in order to pass the carotenoid to its acceptor, Apo-OCP, and this should lead to a limited initial rate of the spectral change, if dimer dissociation precedes the transfer process. The spectrally orange form obtained as the result of the transfer is stable, and its photoconversion is fully reversible (Figure 1B), allowing for an accurate estimation of the photoconversion (O-R) and consequent back-relaxation (R-O) rates (see Figure 1B, and E). The difference between the absorption of the dark-adapted orange and the photoactivated red states (Figure 1A, black line) and the disappearance of vibronic structure upon O-R transition is characteristic for all photoactive OCP species known to date. Analysis of the temperature dependences of the R-O rates of the OCP holoform obtained by carotenoid transfer from COCP revealed ΔH and ΔS values equal to 31.2±1.4 kcal/mol and 56.0±2.3 cal/mo1-K, respectively, which perfectly coincides with the corresponding values for OCP^WT^ determined earlier [14]. Thus, the photocycle of the orange OCP form generated by the carotenoid transfer from COCP is equivalent to the one of OCP obtained by other methods, suggesting that a similar carotenoid transfer process can take place *in vivo* during OCP maturation.

A striking feature of all the time-courses of V-O transitions associated with the carotenoid transfer from COCP to Apo-OCP was a pronounced s-shape, i.e. the rate of the process at the beginning is substantially limited (Figure 1D, black curve). In contrast, changes in O.D. upon the carotenoid transfer from the COCP-W288A mutant (carrying an alanine substitution of the critical Trp-288 [*Synechocystis* OCP^WT^ numbering], which forms an H-bond to the keto-group at the β-ring of the carotenoid) into Apo-OCP were perfectly monoexponential and occurred significantly faster, with no apparent lag-phase. These observations can be readily explained by a higher stability of the wildtype COCP homodimers compared to those of the COCP-W288A mutant, in which the important H-bond between the keto-group of CAN and tryptophan is disrupted [14], which should contribute to destabilization of the dimeric structure of COCP-W288A. Indeed, the difference in the activation energies of the V-O transition for COCP and its W288A mutant is approximately 13 kcal/mol (Figure 1E, Table 1), which we tentatively assign to the difference of two N-H⋯O=C type H-bonds between the keto-group of the carotenoid and the imino nitrogen of the Trp-288 side chain, and this roughly corresponds to other reported values (1 H-bond ~ 5 kcal/mol [36]). Thus, destabilization of the COCP–carotenoid complex owing to removal of the H-bond donors could be correlated with the faster rate of the carotenoid transfer and the absence of the lag-phase at the beginning of the V-O transition (Figure 1D). In turn, this indicates that dissociation of the otherwise very stable COCP dimer may be (i) the reason for the higher energy barrier and (ii) the cause for the initial delay (lag-phase) observed during carotenoid transfer from the wildtype COCP into Apo-OCP. Analogous s-shaped kinetics of OCP-related transitions were also observed during the fluorescence recovery after OCP-induced quenching of phycobilisomes *in vivo*, during which monomerization of FRP occurs and causes an increase in fluorescence recovery rate [14, 37]. Considering the aforementioned facts, we assume that monomerization of COCP should occur first in order to initiate the transfer of the carotenoid molecule.

Remarkably, we observed an extremely low carotenoid transfer efficiency (formation of OCP^O^-like spectral features) in experiments with the Apo-OCP mutant form lacking the first 20 amino acids of the N-terminal extension (NTE) including the αA-helix of the N-terminal domain that normally forms contacts with the C-terminal domain. The NTE is important for stabilization of CTD and NTD interactions [9, 16, 38], and one can suppose that its absence would eliminate steric hindrances in the course of the carotenoid transfer from COCP to OCP. Surprisingly, it appeared that only 5% of Apo-ΔNTE-OCP converted into the orange form upon addition of COCP (Figure 1D), which is an order of magnitude smaller compared to the wildtype Apo-OCP. Thus, the efficiency of carotenoid transfer depends on interactions of Apo-OCP and COCP which require the NTE.

**Table 1.** Thermodynamic characteristics of carotenoid transfer rates (V-O) and rates of the red form back-relaxation (R-O) after the photoactivation. Orange forms were obtained from either Apo-OCP or GFP-OCP^Apo^ chimera upon addition of COCP. Rate constants for V-O and R-O transitions were measured in the range of temperatures from 20 to 45 °C. Values of ΔH and ΔS for transitions were estimated using Eyring-Polanyi equation. Changes of fluorescence intensity of GFP-OCP^Apo^ after addition of COCP was biexponential.

| Carotenoid acceptor upon shuttling | V-O |  |  |  | R-O |  |
| --- | --- | --- | --- | --- | --- | --- |
|  | Apo-OCP | GFP-OCP<sup>Apo</sup> | GFP-OCP<sup>Apo</sup> | GFP-OCP<sup>Apo</sup> | Apo-OCP | GFP-OCP<sup>Apo</sup> |
|  | (absorption) | (absorption) | (fluorescence) | (fluorescence) | (absorption) | (absorption) |
| | | | $\tau_1$ | $\tau_2$ | | |
| $\Delta H$, kcal/mol | $36.8 \pm 1.5$ | $40.2 \pm 4.0$ | $36.2 \pm 3.0$ | $39.6 \pm 2.5$ | $31.2 \pm 1.4$ | $34.9 \pm 1.3$ |
| $\Delta S$, cal/mol·K | $73.4 \pm 2.8$ | $84.5 \pm 7.8$ | $73.9 \pm 5.9$ | $81.3 \pm 4.8$ | $56.0 \pm 2.3$ | $68.6 \pm 2.4$ |

### 3.2. Analytical size-exclusion chromatography (SEC) reveals concentration-dependent dimerization of COCP-W288A and OCP forms with separated domains

The observed unique phenomenon of carotenoid transfer from COCP to Apo-OCP required a careful biochemical characterization of the oligomeric species involved. This analysis was also necessary to account for the fact that not all carotenoids initially coordinated by COCP could finally be stabilized in the orange state upon transfer to Apo-OCP, and we hypothesized, whether some carotenoid-bound side products could be generated in the course of transfer from COCP.

Previously, we demonstrated that, while RCP and Apo-RCP are stable in their monomeric state, COCP shows remarkable propensity to homodimerization [25]. The bound carotenoid stabilized COCP dimers even against manifold dilution, whereas the apoprotein dimers gradually dissociated upon lowering protein concentration [25]. In the present study, by using SEC, we could confirm that COCP-W288A dimers are significantly destabilized due to the mutation of the crucial tryptophan contributing an H-bond to the carotenoid. Even at the highest protein concentration loaded on the column, the SEC profile revealed a mixture of dimers and monomers of the protein, whereas at low protein concentrations, the monomers were prevailing even despite the presence of carotenoid (Figure 2A). Intriguingly, in contrast to stable COCP dimers (Figure 2B), SEC profiles of COCP-W288A followed by carotenoid-specific absorbance at 540 nm at *low* protein concentrations showed the existence of CTD monomers binding carotenoid (Figure 2A), implying the as yet unexplored possibility of carotenoid sliding deeply into the CTD in order to protect the hydrophobic carotenoid from the polar solvent, similar to what is known for RCP [19]. The concentration-dependent behavior of COCP-W288A is similar to that of Apo-COCP (Figure 2A) and supports the idea that the fast kinetics of carotenoid transfer to Apo-OCP (Figure 1D) correlates well with the facilitated dissociation of COCP mutant dimers, and that the W288A mutation entails crucial destabilization of protein-chromophore interactions within the CTD of OCP.

**Figure 2.**
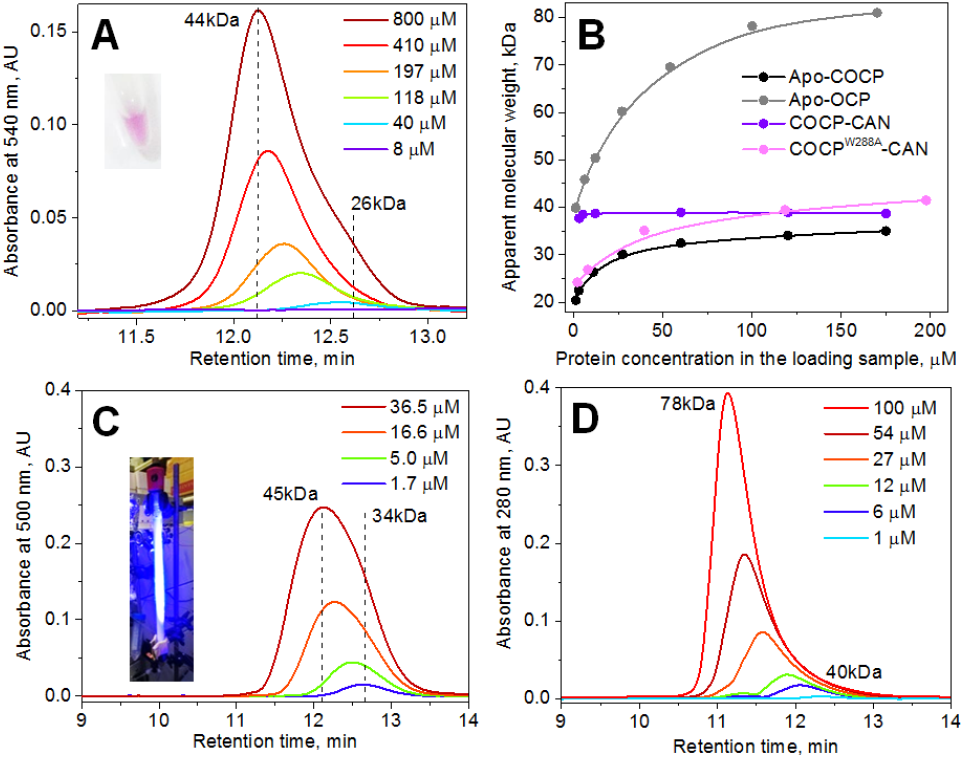
Oligomeric status of COCP-W288A, OCP^R^ and Apo-OCP proteins studied by analytical SEC on a Superdex 200 Increase column. **(A)** – SEC profiles of COCP-W288A (inset) obtained at different protein concentrations loaded on a column and followed by carotenoid-specific absorbance. **(B)** – Dependencies of the apparent molecular weight of different OCP-related species on protein concentration in the loaded sample. **(C)** – SEC profiles of the OCP^WT^ sample pre-illuminated on ice and loaded at different protein concentrations on a constantly blue-LED illuminated SEC column. **(D)** – SEC profiles for different protein concentrations of Apo-OCP followed by protein-specific absorbance. Flow rate was 1.2 ml/min, temperature was 23 °C. The results were reproduced at least two times for each case.

This is in line with our previous results obtained during studies on the purple W288A mutant of full-length *Synechocystis* OCP, which recapitulated structural and functional properties of the photoactivated OCP^R^ form with separated protein domains [14] and also displayed the pronounced concentration-dependent homodimerization [13]. Of note, this is somewhat contradictory to the earlier interpretation by Blankenship and co-workers that OCP photoactivation is accompanied by dissociation of stable OCP^O^ dimers [39], and to the crystal structures, in which OCP^O^ consistently forms a compact dimer [9, 19, 40] suggesting that the real situation of the oligomeric status of OCP is more complex. Our earlier SEC studies revealed that OCP^O^ forms monomers with lesser tendency of self-association than the purple OCP-W288A mutant or Apo-OCP [13, 25, 27]. Questioning whether concentration-dependent dimerization is inherent to any OCP form with separated domains, we analyzed different concentrations of either OCP^R^ or Apo-OCP by SEC (Figure 2C and D). In the first case, the OCP^WT^ sample was pre-incubated on ice and exposed intense blue light illumination, which led to a stable visual photoactivation, and the sample was then loaded on a SEC column operated under strong actinic light (constant blue-LED illumination) to ensure maximal exposure and photoactivation of the sample (Figure 2C). This experiment showed that, even though most likely not the whole population of OCP was converted to OCP^R^, the peak gradually shifted toward earlier elution times, again indicating concentration-dependent protein self-association. This further supports our earlier conclusions that the OCP-W288A mutant mimics properties of photoactivated OCP^R^ [14]. Remarkably, the OCP apoprotein also demonstrated strong concentration-dependent ability to form dimers, suggesting the same mechanism as in the case of OCP^R^ and OCP-W288A and raising the question about which contribution of the OCP protein part is responsible for such a concentration-dependent behavior.

### 3.3. SAXS-based modeling of the OCP and Apo-OCP structures in solution

The fact that in the absence of carotenoid Apo-COCP and Apo-OCP show the same pattern of dimerization (Figure 2B), whereas neither RCP or Apo-RCP dimerized under similar conditions [25], suggests that the most likely mechanism of dimerization of OCP forms with separated domains involves interaction via their CTDs, i.e. through tentative formation of a protein interface expected for COCP dimers. In order to get structural insight and validate our SEC data and interpretations relevant for the carotenoid transfer mechanism, we studied Apo-OCP and OCP^O^ samples by small-angle X-ray scattering (SAXS) and attempted to model their solution structure.

At high protein concentration (200 μM) loaded on a Superdex 200 Increase column coupled to a multiparametric detection system and synchrotron SAXS (Figure 3A), Apo-OCP revealed an asymmetric peak with a skewed Mw distribution, suggesting a mixture of protein dimers (left part of the peak) and their partially dissociated forms (right part of the peak), in agreement with Figure 2D. The scaling and averaging of the SAXS frames from the extreme left part of the peak, corresponding to predominantly dimeric Apo-OCP species, resulted in the curve that could not be approximated reasonably well, neither by a crystallographic OCP^O^ monomer (PDB ID 4XB5) nor by a dimer (PDB ID 3MG1), whose sizes were clearly smaller than suggested by the SAXS curve (R_g_ = 37.3 Å, D_max_ = 170 Å; Figure 3B and C), and, therefore, resulted in very large discrepancies (χ^2^>8). At the same time, if we considered that Apo-OCP is a dimer with separated NTD and CTD domains, in which CTDs are connected within the tentative COCP-like dimeric core (Figure 3C, orange), suggested by SEC analysis (Figure 2) and modeled using CORAL [35] considering the flexibility of the NTD-CTD linkers to minimize the discrepancy with the data, we could obtain a reasonable fit to the experimental SAXS curve (χ^2^=1.2, Figure 3B).

**Figure 3.**
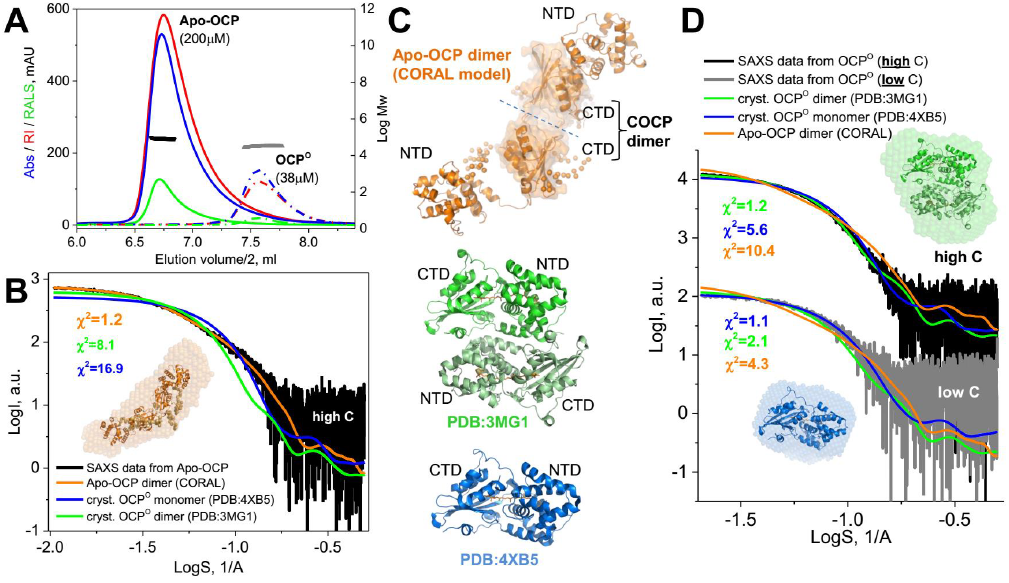
Analysis of Apo-OCP and OCP^O^ by SAXS. **(A)** – SEC profiles of Apo-OCP (200 μM) and OCP^O^ (38 μM) monitored by triple detector array (TDA) consisting in absorbance, refractive index, or right angle light scattering detectors. The flow (0.5 ml/min) was split in two for TDA and SAXS detection which is reflected in halved elution volumes shown on X axis. Temperature was 20 °C. Black and grey thick lines represent Mw distributions over the Apo-OCP and OCP peaks, respectively. **(B)** – SAXS curve (black) corresponding to the extreme left part of the Apo-OCP peak presented on panel A with fits from crystallographic OCP^O^ monomer, OCP^O^ dimer, and a CORAL-derived model of the Apo-OCP dimer (see Materials and methods for further details). Inset, the resulting structural model of the Apo-OCP dimer superimposed with the corresponding *ab initio* envelope from DAMMIF/DAMAVER procedure. **(C)** – corresponding structural models drawn using PyMol 1.6.9. **(D)** – Approximation of the SAXS data for OCP^O^ obtained at high (black) or low (grey) protein concentration by the structural models presented on panel C. Inset, models of the OCP^O^ monomer and dimer superimposed with the corresponding *ab initio* envelopes from DAMMIF/DAMAVER procedure. Color coding is preserved throughout panels B, C, D. Superposition of models with *ab initio* envelopes were made in UCSF Chimera v.1.11 using ‘fit to map’ tool.

Surprisingly, only the SAXS curve obtained at low OCP^O^ concentration (Rg = 22.7 Å, D_max_ = 68 Å; Figure 3D, grey) could be well approximated with the crystallographic OCP^O^ monomer (χ^2^=1.1), whereas a fit of this monomer to the SAXS data recorded at high OCP^O^ concentration (R_g_ = 27.3 Å, D_max_ = 95 Å; Figure 3D, black) was not satisfactory (χ^2^=5.6). Conversely, the crystallographic OCP^O^ dimer could not describe the SAXS data at low OCP^O^ concentration (χ^2^=2.1), but was reasonably well fitting the SAXS curve for the high OCP^O^ concentration (χ^2^=1.2). Importantly, the CORAL-derived model of the Apo-OCP dimer (Figure 3C, orange) could not fit the OCP^O^ SAXS curve obtained, neither at low (χ^2^=4.3) nor at high (χ^2^=10.4) concentration (Figure 3D), strongly suggesting that the concentration-induced dimerization mechanisms of OCP forms with separated and compact NTD and CTD domains may significantly differ. Considering similar features of Apo-OCP, OCP^R^, and OCP-W288A such as the increased hydrodynamic size and concentration-dependent behavior, we assume that the SAXS-based model of the Apo-OCP dimer is principally applicable to the photoactivated OCP or its analog. At the same time, we cannot exclude that these forms with separated domains can generate also other dimeric assemblies upon increasing protein concentration, e.g. stabilized by NTD-CTD interactions, however, in the absence of stabilizing carotenoid, i.e. in Apo-OCP, these interactions seem less probable. Since the formation of the Apo-OCP-like dimer at high protein concentrations decreases the rate of R-O relaxation (SI) we propose that such structures may not only be a feature of *in vitro* experiments, but may also play a certain role in regulation of cyanobacterial photoprotection.

Thus, the oligomeric state of both donors of carotenoids (COCPs) and acceptors (Apo-OCP) is important for the carotenoid transfer and cannot be disregarded.

### 3.4. Carotenoid transfer evidenced from biochemical studies

A critical point needing clarification is the limited efficiency of OCP^O^ formation in the course of carotenoid transfer from COCP into Apo-OCP, since some purple-violet forms always remained present even in Apo-OCP excess as shown in Figure 1A. The concentration-dependent assembly patterns of the various OCP and COCP species (Figure 2 and 3) suggest that the outcome of mixing experiments could include a variety of spectral and structural species, which may have included not only some remaining COCP (which by some reason does not interact with Apo-OCP), but also due to the stabilization of the carotenoid between CTDs of other OCP species or other CTD-CTD interactions. Therefore, we investigated the outcome of mixing experiments as shown in Figure 1 by analytical SEC and gel electrophoresis (Figure 4). Indeed, Figure 4A shows that the ~32 kDa orange OCP is not the only product of the carotenoid transfer, but, in addition, there is a fraction of a heavy ~69 kDa carotenoid-containing species (Figure 4B). The SDS-PAGE of this fraction shows that these species consist exclusively of full-length (Apo)-OCP with calculated Mw of 36 kDa (Figure 4B). Strikingly, the absorption of this heavy fraction represents a mixture of orange and violet species and was (at least partially) photoactive (Figure 4C). Considering the fact that the carotenoid should interact with both, one NTD and one CTD, in order to achieve the orange spectrum and maintain photoactivity, we assume that the orange 69 kDa dimers are the result of cross-domain binding of one carotenoid between two former Apo-OCPs. However, the spectral results suggest two different arrangements: In the orange species, the carotenoid is coordinated between NTD and CTD of two different Apo-OCPs, whereas in the violet species, the carotenoid links the CTDs of two Apo-OCPs.

**Figure 4.**
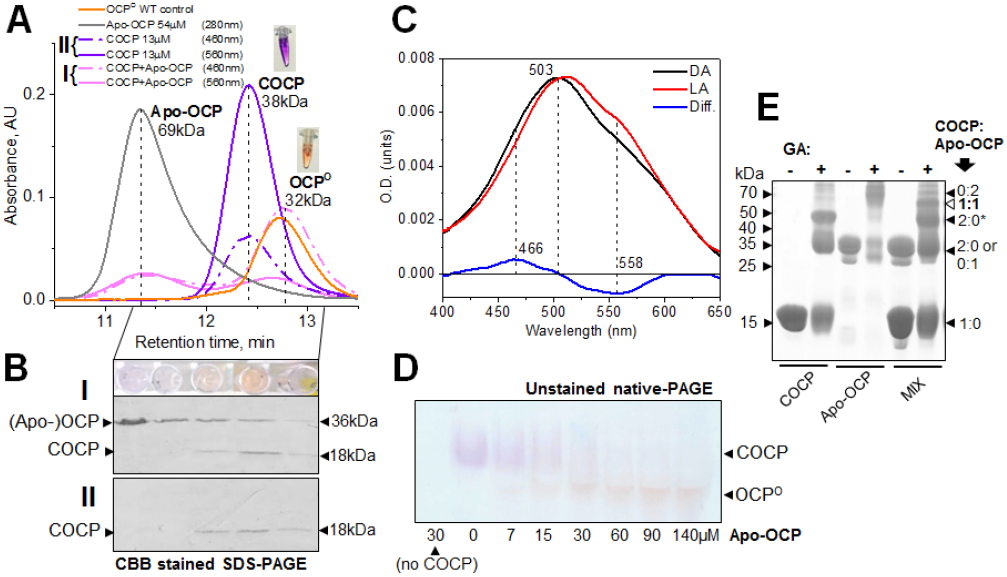
Carotenoid transfer followed by SEC and native gel-electrophoresis. **(A)** – SEC profiles of COCP, Apo-OCP, and of products of carotenoid transfer obtained by mixing COCP and Apo-OCP monitored by absorbance at indicated wavelengths. Note that dual wavelength detection allows revealing of the spectral shift upon carotenoid transfer accompanying formation of OCP^O^, whereas the fraction at ~11.2 min has almost equal absorption at 460 and 560 nm. OCP^O^ sample was loaded as the control. Inset shows the color of COCP (donor of carotenoid) and OCP^O^ (product of transfer). Dashed lines show positions of the corresponding maxima of the peaks of Apo-OCP, COCP, and OCP^O^. **(B)** – SDS-PAGE analysis of the fraction obtained from the COCP profile (II) or its mixture with Apo-OCP after the completion of the carotenoid transfer (I). Fractions of I profile are shown above the gel. SDS gels were stained by Coomassie brilliant blue (CBB). Positions of protein bands and those of Mw markers are indicated. **(C)** – The absorption spectra of the fraction collected at ~11.2 min in dark-adapted (DA) or light-adapted (LA) states and their difference spectrum showing some photoactivity. Dashed lines show characteristic spectral features. **(D)** – Carotenoid transfer followed by unstained native gel-electrophoresis. COCP was mixed with increasing amounts of Apo-OCP, incubated for 30 min at 33 °C and then loaded on the gel. Controls did not contain either COCP (first lane) or Apo-OCP (second lane). Positions of carotenoid-containing proteins are indicated by arrows. **(E)** – COCP (20 μM), Apo-OCP (40 μM), or their mixture were pre-incubated for 30 min at room temperature and then mixed with glutaraldehyde (GA – added to only even lanes) for chemical crosslinking of the carotenoid transfer intermediates. Mw standards are indicated to the left (in kDa). To the right are indicated determined oligomeric states of the crosslinked species formed (number of COCP subunits:number of Apo-OCP subunits). Asterisk marks the band, presumably corresponding to alternatively crosslinked COCP dimers (SEC does not support COCP oligomers above dimers). Empty triangle shows the ~55 kDa band most likely corresponding to the 1:1 hybrid COCP:Apo-OCP complexes, i.e., tentative intermediates of the carotenoid transfer.

We were also able to visualize the carotenoid transfer and to physically separate the violet COCP (donor) and orange OCP^O^ (the product of the transfer to Apo-OCP) species by gel-electrophoresis under non-denaturing conditions (Figure 4D). Remarkably, this did not require any staining and resulted in a clear transition of the COCP into OCP^O^ band. Interestingly, the latter could be photoactivated upon illumination of the gel by blue-LED causing an apparent “redding” of the bands (not shown).

Moreover, by using chemical crosslinking of COCP/Apo-OCP mixtures in the course of the carotenoid transfer (Figure 4E) we could reveal formation of the tentative intermediary hybrid COCP:Apo-OCP species (55 kDa), which were not detectable by SEC after the completion of the carotenoid transfer and were also not present when the individual COCP or Apo-OCP underwent crosslinking (Figure 4E). Formation of the crosslinked COCP:Apo-OCP band (band “1:1”) inhibited Apo-OCP dimerization (band “0:2”), in line with the assumed CTD-CTD contacts formed within such Apo-OCP dimers and the proposed SAXS-based model (Figure 3C, orange). Intriguingly, crosslinking of COCP resulted in a more complex pattern than was expected from its 38 kDa dimers detected on SEC [25]. Indeed, two new almost equally intensive bands having electrophoretic mobility corresponding to 37 and 46 kDa, besides the 18 kDa band corresponding to COCP monomer, could be found on the gel. Since SEC did not support the presence of COCP trimers (Figure 2B) and because the intensity of the 46 kDa band was not weaker than that of the 37 kDa band (expected for crosslinked trimeric and dimeric protein species, respectively), we speculate that these two bands correspond to differently crosslinked COCP polypeptide chains.

### 3.5. Fluorescence of GFP-OCP chimera reveals intermediates of the carotenoid transfer

As outlined above, the mechanism of carotenoid transfer from COCP into Apo-OCP is a multi-step process including a series of biochemical intermediates, from which not all are endowed with distinct absorption spectroscopic features. In order to identify more details of the transfer process by spectroscopy, we used a different strategy to monitor the actual position of the carotenoid based on fluorescence resonance energy transfer from an additional chromophore attached to OCP. Previously, we used fluorescent dyes to study the OCP photocycle [15, 20, 27]. This approach is based on measurements of excitation energy transfer (EET) from some exogenously introduced fluorophore to the carotenoid of OCP. However, application of organic dyes has several significant shortcomings such as non-specific binding, multiple donors per acceptor of energy, difficulties associated with obtaining an adequate model of the donor species in the absence of an acceptor, etc., which makes such a system difficult for evaluation of protein conformational changes. Thus, for this specific study we decided to introduce the 28.5 kDa enhanced green fluorescent protein (further GFP) at the N-terminus of OCP close to the αA-helix. GFP was placed in a close proximity to the NTE, as experiments with the ΔNTE deletion construct revealed that this part of OCP is important for carotenoid transfer.

**Figure. 5.**
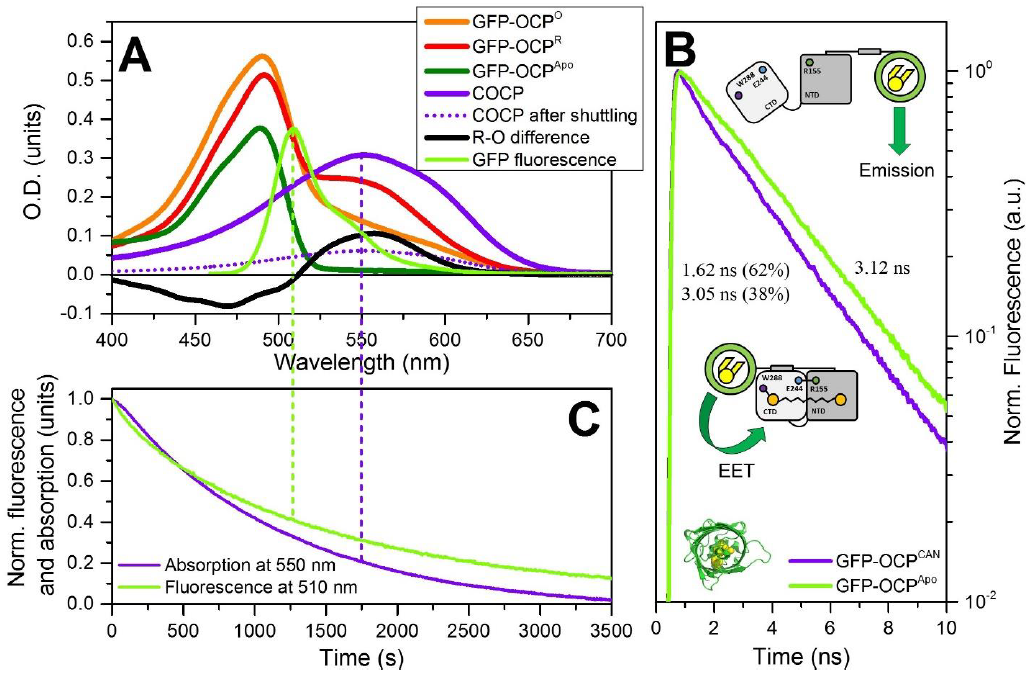
Carotenoid transfer from COCP to GFP-OCP^Apo^ chimera. **(A)** Absorption spectra of GFP-OCP chimera and related species. Upon addition of 4.4 μM of COCP (violet) to 6.3 μM solution of GFP-OCP^Apo^ (green) absorption of COCP gradually decreases. After equilibration of COCP - GFP-OCP^Apo^ interactions resulting spectrum of the system (orange line) represents sum of GFP, OCP^O^ and COCP absorption (violet dashed line). Obtained orange fraction is photoactive and, upon illumination of the sample by actinic light (450 nm, 200 mW), reversibly converts to the red state (red line). Difference (black line) between the spectra of red and orange states is typical for all known OCP species. **(B)** – GFP fluorescence decay kinetics of GFP-OCP chimera in the absence (GFP-OCP^Apo^) and in presence of carotenoid (GFP-OCP^CAN^). COCP to GFP-OCP^Apo^ ratio was equal to 3. Insets show the structure of GFP and schematic representation of GFP-OCP chimera. **(C)** – kinetics of carotenoid transfer monitored by measurements of O.D. at 550 nm and intensity of GFP fluorescence at 510 nm, simultaneously. Experiment was conducted at 20 °C and constant stirring.

**Figure 5.**
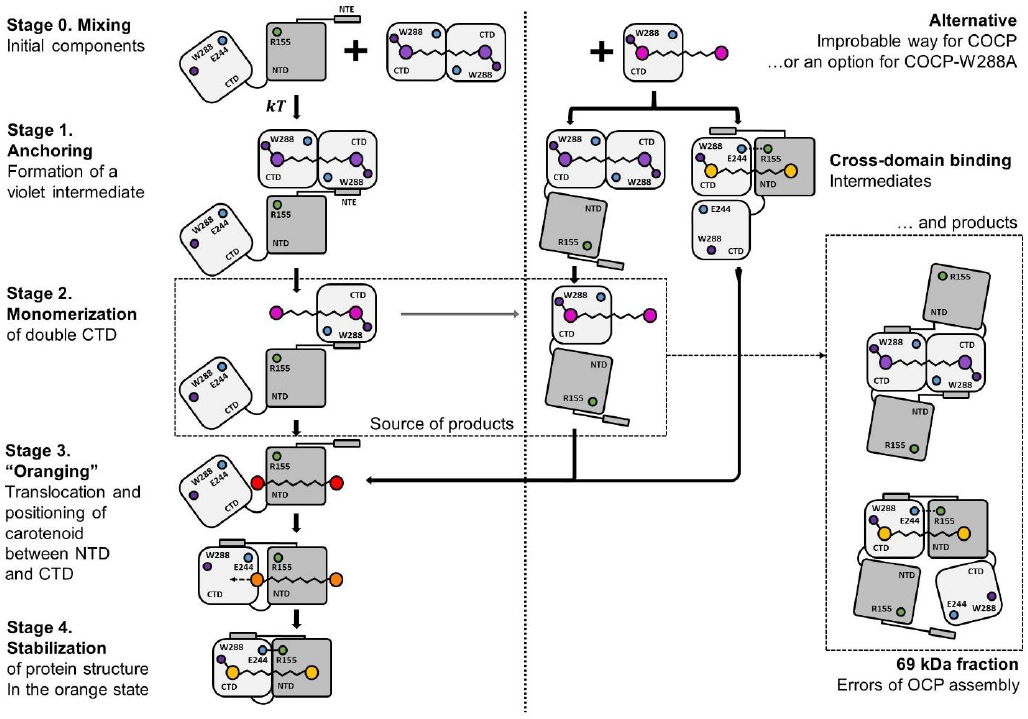
Working model of the carotenoid transfer from COCP to Apo-OCP leading to reconstruction of a photoactive OCP. After mixing (stage 0), COCP is anchored by Apo-OCP (presumably, via NTE) (stage 1) and undergoes monomerization (critical stage 2) in order to transfer carotenoid into the NTD of Apo-OCP. Since the NTD has higher affinity to the carotenoid molecule than the CTD, it accepts the carotenoid from one of the CTD subunits of the anchored COCP. This leads to closure of OCP with bound carotenoid into the compact OCP^O^-like structure (stage 3) stabilized by carotenoid (stage 4). Efficiency of carotenoid transfer is high, and over 70% of carotenoid from COCP is transmitted to the orange photoactive form. However, as Apo-OCP tends to form homodimers at high concentrations, such structures could be stabilized by cross-domain carotenoid binding (confirmed by the data on Figure 4). Alternative pathway (right part) requires preliminary monomerization of COCP (effectively achieved by COCP-W288A mutant) or involvement of some other carotenoid carrier.

The absorption spectrum of the GFP-OCP^Apo^ chimera in the visible region is related exclusively to the absorption of GFP (Figure 5A) with a maximum at 488 nm. Subsequent mixing of COCP and GFP-OCP^Apo^ results in significant changes of absorption accompanying the formation of the orange state, equivalent to such transitions described for Apo-OCP (see Figure 1). After introduction of the carotenoid into GFP-OCP^Apo^ and formation of the photoactive orange state, we observed an appreciable decrease in GFP fluorescence lifetime. The major component (62%) of the fluorescence decay was characterized by a lifetime of 1.62 ns, while in the initial GFP-OCP^Apo^ the decay was monoexponential with a lifetime of 3.05 ns (Figure 5B). EET efficiency was estimated to be equal to 42.5 %. We calculated the overlap integrals and corresponding Förster radii – 56.9 Å and 59.6 Å for OCP^O^ and COCP, respectively. The observed EET efficiency corresponds to a distance between the donor and acceptor of about 61 Å, which is approximately the size of the OCP molecule. This observation indicates that the orientation factor к^2^ could be low (transition dipoles are close to perpendicular), since the real distance between the carotenoid and the GFP chromophore is definitely less than 60 Å. Unfortunately, no significant difference between the lifetimes of GFP-OCP in its orange and red state was found experimentally, which is probably due to small changes of the donor-acceptor distance or the unfavorable orientation factor. However, we can conclude that GFP fluorescence is sensitive to the presence of carotenoid in close proximity, thus, the fluorescence of GFP allowed us to study carotenoid transfer, as its products are characterized by EET to the carotenoid cofactor, which is absent in GFP-OCP^Apo^.

We studied the kinetics of either absorbance or GFP fluorescence changes during carotenoid transfer upon mixing of GFP-OCP^Apo^ and COCP, and the most striking differences were found in the shape of the respective time-courses, especially their initial parts (Figure 5C). Changes of GFP fluorescence occurred without any lag phase and could be perfectly approximated by the sum of two exponential decay exponents. The fast component of GFP fluorescence quenching upon the transfer is significantly faster than the changes of 550 nm absorption measured under exactly the same experimental conditions. Thus, GFP seems to be sensitive to the formation of an intermediary complex with a violet carotenoid in the CTD-CTD arrangement, even before the carotenoid gets into its final position between the CTD and NTD. Analysis of the temperature dependencies of the rates of absorption changes and of the rates of fluorescence decay components revealed that the slow component of GFP fluorescence decay and the changes in absorption have the same ΔH and ΔS values (*Table 1*), strongly suggesting that they are related to the same processes. The slow component of fluorescence changes is probably associated with the adjustments of protein structure that are following relocation of the carotenoid and are necessary for stabilization of the orange form. Thus, carotenoid transfer is initiated by formation of an intermediate complex with a carotenoid carrier, violet COCP, where changes of carotenoid absorption did not occur yet. We may refer to this phase of carotenoid transfer as the “anchoring stage”.

## Conclusions

We want to summarize the results of our study by proposing a working model of the carotenoid transfer mechanism between COCP and Apo-OCP (Figure 5), which might occur during chromophore insertion and assembly of photoactive and photoprotective OCP *in vivo*. It is safe to assume that incorporation of the carotenoid should not occur co-translationally, but requires synthesis and folding of the OCP apoprotein. However, carotenoids are synthesized, processed and stored at or in membranes, while Apo-OCP is a water-soluble protein, thus the mechanism of carotenoid transfer through the aqueous cytoplasm is not clear. One could assume that an Apo-OCP, with its open domain structure, could drag a carotenoid out of the membrane by itself, however, the presence of water-soluble carotenoid carriers such as COCPs could significantly facilitate the assembly of holo-OCP and play important role in regulation of the cellular holo-OCP content for photoprotection in cyanobacteria. Of note, many cyanobacterial species harbor one or several separate genes for CTD and NTD (HCP) analogs [23, 24]. Whereas the physiological role of NTD (HCP) analogs is well established, the as yet orphan CTD homologs could be assigned a physiological role at least during maturation and regulation of fully functional OCPs cycling.

In this work, we show how the simple addition of OCP apoprotein readily leads to the breaking up of the otherwise very stable COCP dimer, eventually extracting the carotenoid from this assembly, which involves carotenoid translocation from one hydrophobic protein environment into another thus obviating a passage through the solvent. To the best of our knowledge, this is the first description of a protein-to-protein carotenoid transfer. In this unprecedented OCP case, the process predominantly results in the formation of the orange OCP form, which undoubtedly represents OCP^O^ in the monomeric state (see Figures 1 and 2). However, our results also suggest that formation of OCP^O^ upon carotenoid transfer passes through several different and spectroscopically distinguishable intermediate states. First of all, the NTE, which is responsible for stabilization of CTD-NTD interactions in the orange form, may bind COCP (Figure 5, Stage 1), which, being a dimer of two OCP-CTDs, has two sites for interactions with the NTE. Binding of COCP via NTE increases the probability that Apo-OCP can approach the carotenoid-containing CTD, when spontaneous monomerization of COCP due to protein dynamics [25] may occur (Figure 5, Stage 2). The existence of such a violet intermediate was demonstrated by fusion of GFP to the N-terminus of OCP (see Figure 4), which appeared to be sensitive to the presence of carotenoid. We may hypothesize that the interaction with the NTE may also affect the stability of COCP and initiate its monomerization, which requires disruption of a critical hydrogen bond between the keto-oxygen of carotenoid and Trp-288 in one of CTDs. Subsequently, the carotenoid enters the NTD of Apo-OCP, a process, which is determined by its higher affinity to NTD that is supported by the fact that carotenoid transfer from the NTD into Apo-CTD could not be observed. After this crucial transfer step, stabilization of the orange state occurs as a regular and well-described relaxation of the red form. At this stage (3) the most significant changes of carotenoid absorption could be monitored. We assume that the rate of carotenoid transfer depends on the characteristic rate of R-O relaxation (as a limiting step) and the concentration of COCP monomers (or Apo-OCP–COCP complexes).

In fact, the absence of the NTE dramatically reduces carotenoid transfer efficiency (see Figure 1D), though not abolishing it completely, which indicates that the NTE-CTD interaction may be involved in COCP recruiting or anchoring by Apo-OCP. Alternative ways for transfer require the carotenoid carrier to be in the monomeric state (which is a minor state for COCP, but a major state for its W288A mutant, see Figure 2). Of note, several reaction intermediates could be obtained as side products of the proper carotenoid transfer process due to the ability of two CTDs to form a homodimer, and NTD and CTD to form heterodimer, which are stabilized by carotenoid. This way is not effective if the initial carotenoid carrier is as stable as COCP, however, destabilization of COCP (e.g., by W288A mutation) leads to a significant increase of the transfer rate. The question of carotenoid content in the CTD carrier is also important, as *in vivo* OCP binds echinenone (ECN) or 3’-hydroxy-ECN, thus stabilization of CTDs protein-protein interactions by such mono-ketolated carotenoids by two H-bonds is impossible, thus leaving unketolated (hydroxylated) carotenoid β-ring exposed for possible interactions with the NTD of Apo-OCP. Thus, our *in vitro* experiments revealed several hidden features of OCP assembly and highlights a particular role of the NTE, which not only stabilizes CTD and NTD interactions in holo-OCP, but also plays important role in the initial stage of Apo-OCP-COCP interaction to greatly facilitate the reconstruction of the photoactive OCP from its apoprotein form. In the cell, COCP and probably its individual homologs could serve as carotenoid depot that readily supply their cargo to full-length OCP when the cell turns on OCP synthesis for photoprotection.

## Acknowledgements

N.N.S. is grateful to Dr. Cy Jeffries (EMBL Hamburg) for his help during SAXS data collection, reduction and initial processing. This work was supported by grants from Russian Foundation for Basic Research 15-04-01930a to E.G.M. E.G.M. was supported by Dynasty Foundation Fellowship. The reported study was funded by RFBR and Moscow city Government according to the research project 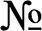 15-34-70007 ≪mol_a_mos≫. T.F. acknowledges the support by the German Federal Ministry of Education and Research (WTZ-RUS grant 01DJ15007) and the German Research Foundation (Cluster of Excellence “Unifying Concepts in Catalysis”).

## Supplementary information

Additionally, we have noticed that R-O conversion rates are significantly slower in the sample which initially had carotenoid in COCP-W288A. This effect could be explained by the presence of significant amount of Apo-COCP-W288A in the sample, which probably may interact and form intermediates with the red form stabilizing it and preventing relaxation of to the red form. In order to test this hypothesis, we studied dependency of R-O rate on concentration of Apo-COCP (CTD). Upon the increase of Apo-COCP content we observed a gradual decrease of R-O rates which proves that CTDs can interact with the red form of OCP.

**Figure S1.**
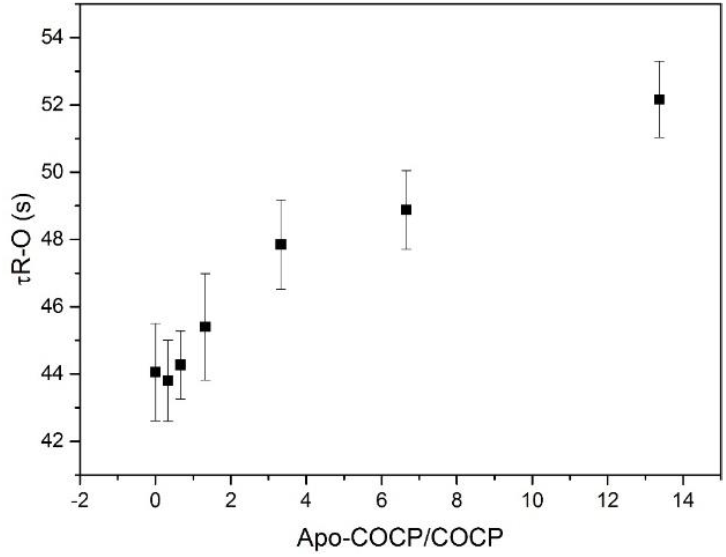
Dependency of time constants of red to orange relaxation of reconstructed OCP on concentration of Apo-COCP. Orange form was obtained by mixing COCP and Apo-OCP in 1:2 ratio. Experiments were conducted at 33 °C.

